# Characterising the Behaviour of the Ctenophore *Pleurobrachia pileus* in a Laboratory Aquaculture System

**DOI:** 10.1101/2020.05.25.114744

**Authors:** Amy Courtney, George O.T. Merces, Mark Pickering

**Affiliations:** School of Medicine, University College Dublin, Ireland; UCD Centre for Biomedical Engineering, University College Dublin, Ireland

**Keywords:** ctenophore, husbandry, model organism, evolution, behaviour

## Abstract

Neurobiological research focuses on a small number of model organisms, broadening the pool of animals used in research may lead to important insights into the evolution of nervous systems. The ctenophore is emerging as a promising model, but we are currently lacking an understanding into the relationship between behaviour and environment which is in part due to a lack of a standardised long-term laboratory husbandry system. We established a collection and husbandry system for wild caught *Pleurobrachia pileus*. We examined the behavioural profile of the animals over time in this controlled environment. We could reliably catch them on a seasonal basis, and we could keep the animals alive in our specialised aquarium system for months at a time. *P. pileus* spends most of the time in an inactive ‘drifting’ state which is interspersed with periods of one of 5 active behaviours. The most common active behaviours are tentacle resetting and feeding. The longest duration behaviours include swimming up or down. Time of day does not appear to alter their behavioural profile. Gaining a better understanding of the behaviour of these animals has important implications for systems and evolutionary neuroscience.

## 1 Introduction

Traditionally, the term “model organism” referred to an animal that was “simpler” than humans that was amenable to experimentation and facilitated investigation of a specific biological process. The meaning of the term has changed in recent years and is now often used to describe an organism which is utilised by many researchers and for which a significant experimental toolbox is available. Thus, originally model organisms were selected for suitability and convenience while now their selection is based on resource availability [Russell et al., 2017]. The catalogue of life [Roskov Y. et al., 2019] currently lists ~1.3 million animal species and it has been predicted that the true value is closer to ~7.7 million [Moraet al., 2011]. Using neuroscience as an example, 75% of research focuses on rat, mouse and human brains with invertebrates accounting for 7.9% [Manger et al., 2008]. This is striking when you take into consideration that invertebrates make up 95% of all extant animal life [Lewbart, 2011]. While recent statistics on the prevalence of specific model organisms in neuroscience are not available, it appears that the animals most commonly used include human, rats, mice, fruit flies (*D. melanogaster*), zebrafish (*D. rerio*) and roundworms (*C. elegans*) [Borniger, 2015, Maximino et al., 2015, Chen, 2019]. While focusing research efforts on a small number of organisms has advantages, it is important to acknowledge the limitations and disadvantages that come with this approach which could be overcome by broadening the pool of model organisms used. In addition, the established model organisms may not be appropriate for answering all questions. The Krogh principle states that amongst the diversity of life there is one or multiple organisms which is most appropriate for answering a specific physiological question [Krogh, 1929]. To that end, we need to expand our repertoire of model organisms to include animals at specific positions on the phylogenetic tree while also taking into consideration the new opportunities they may present to answer specific neurobiological questions.

Members of the phylum Ctenophora are emerging as promising model organisms for comparative neurobiological investigations. Their position on the phylogenetic tree is contentious with some studies placing them as the sister taxon to all animals, while others support the sponge-first scenario [Dunn et al., 2008, Ryan et al., 2013, Feuda et al., 2014, Moroz et al., 2014, Borowiec et al., 2015, Pisani et al., 2015, Arcila et al., 2017, King and Rokas, 2017, Simion et al., 2017]. It has also been suggested that these animals may have evolved their nervous system independently to all animals [Moroz et al., 2014], however this claim remains controversial [Marlow and Arendt, 2014, Jékely et al., 2015]. Thus, these animals are either our best example of what the first nervous system looked like, or they represent the only example of divergent nervous system evolution. Gaining a better understanding of these animals will undoubtedly provide interesting insights into the foundational principles of neural structure and function. In Irish waters the most common species of ctenophore is *Pleurobrachia pileus* [Müller, 1776] or the sea gooseberry [Neal, 2005]. Ctenophores possess other features which allude to their promise as a model organism. They are small (0.1mm-20mm) and transparent which facilitates imaging studies, they have relatively fast reproduction rates [Greve, 1970, Baker and Reeve, 1974, Martindale, 1987] which facilitates laboratory rearing and genomic/transcriptomic data is available [Ryan et al., 2013, Moroz et al., 2014] which aids in the development of genetically engineered organisms. From a neurobiological perspective, they appear to possess a structurally simple and tractable nervous system, a nerve net distributed across their body [Hernandez-Nicaise, 1973a, b, Jager et al., 2011]. Despite this ‘simple’ neural architecture they display surprisingly sophisticated behaviours [Tamm, 2014]. However, the behaviours appear to be simple and predictable enough to examine quantitatively. Despite the apparent advantages these animals possess as a model organism, there are many unanswered questions concerning their nervous system.

The nervous system evolved to enable animals to quickly adapt to challenges they faced in their specific ecological niches. Decades of behavioural research in conventional model organisms has led to a deep understanding into the relationship between behaviour and environment. However, for unconventional model organisms, such as *P. pileus*, the relationship between behaviour and environment is less clear. For example, the sensory capabilities of these animals are unclear and in-depth analysis of their behavioural repertoire over time has never been performed. Developing a baseline behavioural profile for *P. pileus* will allow us to assess their amenability as a model organism and may elucidate some of the sensory mechanisms triggering these responses. To achieve this, we need a controlled environment. However, as an unestablished model organism, a standardised longterm husbandry system has never been comprehensively described. In order to promote the use of these animals as a model organism for neurobiology and to understand how their behaviours relate to their umwelt, it is imperative that we comprehensively describe the aquaculture system used to study wild caught *P. pileus*, and the characterise the behaviours displayed in this environment.

## 2 Methods

### 2.1 *Pleurobrachia pileus* Collection

Ctenophores were collected from the Irish Sea primarily from Howth Harbour (53°23’34.9″N 6°03’58.0″W) from September 2015 to August 2019. The highest density of animals occurred from April to December (Figure 1A) and lower densities in January and March, using a combination of collection techniques. Specifically, we used both commercial (NHBS Ltd) and custom-made plankton nets with weighted closed cod ends (1.2 - 2.2L capacity), 960-1000mm in length with 250µm and 500µm mesh size. We also used 2.5mm mesh dip nets and dip cups when animals were at high enough density to be visible in the water. Animals were transported from the collection site in 2L containers, at a maximum density of 150 animals/L. Time from collection to transfer to aquarium was ~3 hours, and temperature was not controlled during this time. Animals were not acclimatised to aquarium water before transfer, but we verified that the salinity in the transportation vessel and the aquarium system was comparable (30±2 ppt).

**Figure 1.**
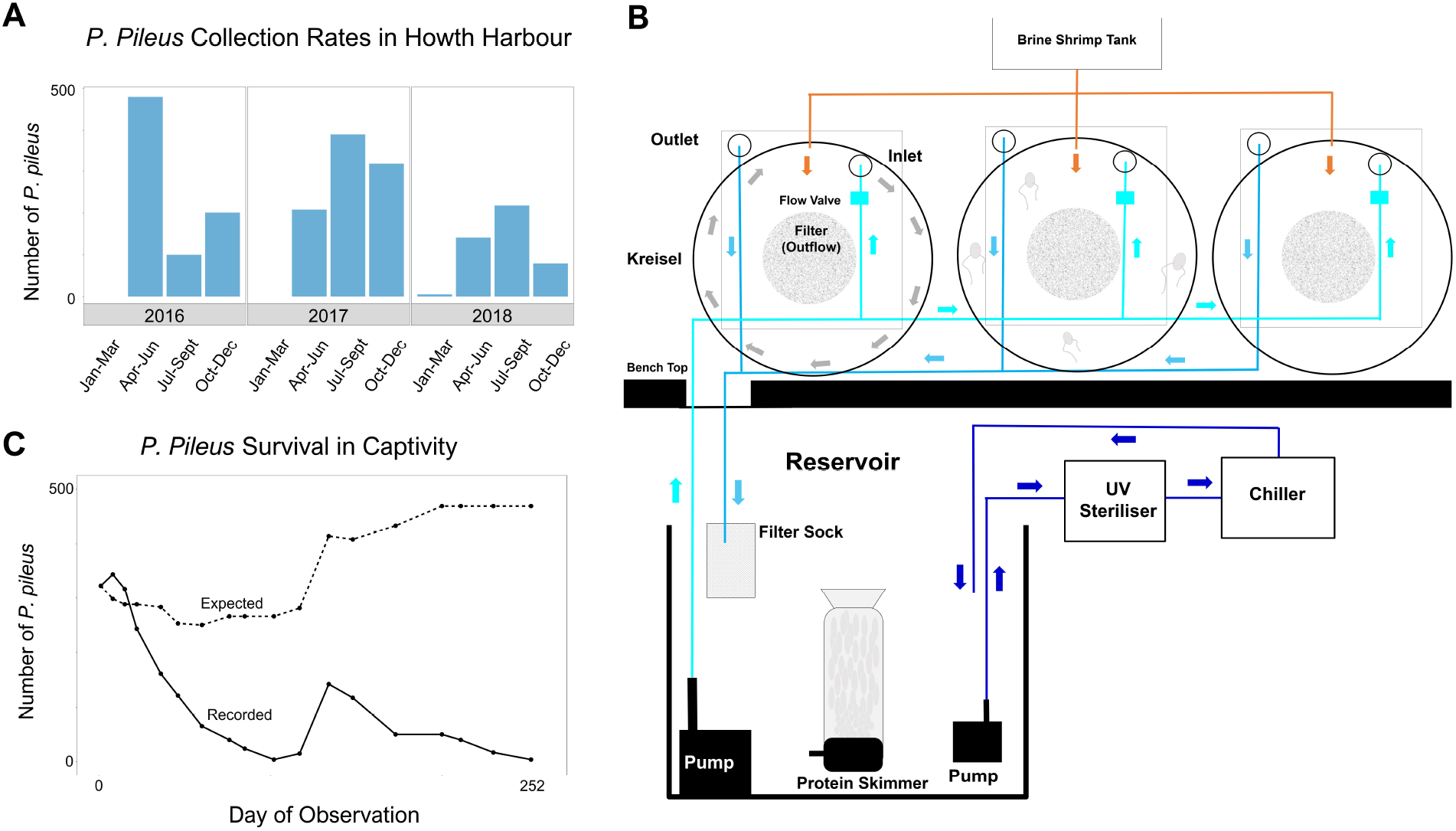
*Pleurobrachia pileus* Collection and Husbandry. (A) Over the course of three years collecting *P. pileus* at Howth harbour we observed a seasonal occurrence of animals, highest densities from April-December and little to no animals in January-March. (B) This is a schematic representation of the husbandry system we established for wild caught *P. pileus*. A parts list for all components in this schematic can be found in Supplementary Table 1. Animals were maintained in kreisel tanks which have a circulating flow which maintains the animals suspended in the water column. Water flows in through the spray bar and is drained through the filter at the back of each tank. An Arduino controlled pump system delivers Artemia nauplii from a holding tank at 20-minute intervals. Water drains from the kreisel tanks to the reservoir initially passing through a filter sock. Live rock, filter balls, a protein skimmer and carbon are present in the reservoir and provide filtration. The water is then pumped into a UV steriliser and subsequently into a chiller (set to 9 ◦C). The cooled water then recirculates back into the reservoir and is subsequently pumped back into the kreisel tanks. Each kreisel possesses a valve to adjust flow. The system has a 150L capacity and we use fresh seawater. (C) The recorded number of animals present in our system compared to the expected number. The expected values relate to the predicted number of animals present assuming no death while also taking into account the number of animals removed for experiments and added after subsequent collection trips. The discrepancy between the recorded and expected values means that the number of animals is decreasing over time due to death.

### 2.2 Establishing and Maintaining the Aquaculture System

A detailed parts list can be found in supplementary table 1 and each component has a designator which will be referred to throughout this text. We established an aquaculture system for *Pleurobrachia pileus* on a bench surface in a multi-purpose laboratory environment (Figure 1B). Due to this, light dark cycle and room temperature was not tightly controlled. Animals were maintained in kreisel tanks (AC1: diameter: 400mm, depth: 200mm; Schuran Seawater Equipment, Jülich Germany). A circulating flow which maintains the animals suspended in the water column is generated by water entering through the spray bar and draining through the centrally positioned filter at the back of the tank (diameter: 15mm, pore size: 3mm). This white filter was covered with a 400µm black mesh which increased visibility of the largely transparent animals while also maintaining artemia nauplii in the tanks by preventing their removal through the larger filter pores. Artemia nauplii (AM12) are delivered from a holding tank (AC18) at 20-minute intervals for continuous automated feeding (using an Arduino controlled pump). Water drains from the kreisel tanks to the reservoir (AC13) initially passing through a 100µm mesh size filter sock (AC8). Live rock (AC7) and the filter start solution (AC16) provided the nitrifying bacteria. To increase the surface area upon which these bacteria can colonise filter balls (AC11) were incorporated into the reservoir. Other filtration components include a protein skimmer (AC12) and carbon (AC10) which are present in the reservoir. The water is then pumped (AC4) into a UV steriliser (AC6) and subsequently into a chiller (AC2). The chiller is set to 9◦C however it fluctuates throughout the year depending on the temperature in the room. It usually stabilises around 12◦C however temperatures up to 17◦C were observed on multiple days during the summer months. The cooled water then recirculates back into the reservoir and is subsequently pumped (AC3) back into the kreisel tanks. Flow was maintained at approximately 600ml/min/tank. Flow to each kreisel can be independently controlled or stopped for cleaning or decommissioning. Each kreisel has a ~28L capacity and the capacity of the entire system is usually ~150L. We use fresh seawater (Seahorse Aquariums Ltd.) and a 25L water change is performed every 3 weeks. Cleaning procedures are performed monthly and involves scrubbing the inner surface of the kreisel tanks and spray bar to remove any debris which has accumulated. No chemicals are used during cleaning.

Animals were added to the aquaculture system and removed for experimentation at various time points, allowing us to calculate the expected number of animals in the system at any point in time. The actual number of animals was measured and showed a decline over time (Figure 1C). This suggests that while the aquaculture system may not be optimal for long term survival of *P. pileus* in the laboratory, it is possible to maintain animals for extended periods of time (i.e. several months), which is sufficient for the experimental aims of this study.

### 2.3 Behavioural Monitoring

Animals used for behavioural experiments were caught in October 2018 and had been in the tanks for 18 days before the experiment began. The animals were subjected to a 12-hour light dark cycle for the duration of the experiment. 13 animals were present in one kreisel tank (one animal per 1.9L), and 10 minute videos were acquired of this single tank at the start (7am), mid (1pm ± 1 hour) and end (7pm ± 1 hour) of the light cycle over a four-day period. No data was collected during the dark cycle. The same animal could not be tracked from one video to another. Specific, discrete behavioural states were defined: t he a ctive b ehaviours observed in this environment were tentacle resetting, feeding, swimming up, swimming down, and escape swimming, while drifting was considered their default ‘inactive’ behavioural state (supplementary table 2, supplementary video 1). Videos were manually annotated with Kinovea software [Charmant] with the time and location of transition between behavioural states in each animal. Periods of lost data or unclassifiable behaviours due to environmental interference and motionlessness were also noted. Motionlessness refers to animals coming into close contact with the black mesh on the filter outflow and staying in a fixed position for periods of time. Environmental interference relates to animals with a tentacle attached to the inner surface of the tank or to another animal.

### 2.4 Statistical Analysis and Data Visualisation

The statistical analysis and data manipulation was performed using R [Core Team, 2018]. R packages utilised for data visualisation include ggplot2 [Wickham, 2016], ggthemes [Arnold, 2019], cowplot [Wilke, 2019] and networkD3 [Allaire et al., 2017]. For behavioural analysis, we examined the frequency and duration of individual behaviours, the temporal sequence of behaviours, and the location within the tank in which specific behaviours occurred.

## 3 Results

### 3.1 Behaviour Distributions Across the Light Cycle

The sequence of behaviours of each of the 13 animals over each of the 10 minute recordings at the start, middle and end of the light cycle over 4 days is shown in Figure 2A. Pooled across all recordings, the majority of the time is spent drifting (inactive behaviour); ~74% of the total time analysed in the dataset was spent drifting, ~8% was unclassifiable and ~18% was spent actively behaving. Of that, tentacle resetting accounted for ~9.2%, feeding was ~2.4%, swimming down equated to ~6%, swimming up was ~0.5% while escape swimming was ~0.2%. When this is segregated into the start, middle and end of the light cycle, there is an apparent decrease in the total amount of time spend engaged in active behaviours as the light cycle progresses (Figure 2B). This could result from changes in the duration of behaviours, or changes in the number of times a behaviour is initiated. However, when the total number of behavioural initiations is pooled, there is an apparent increase in initiations as the light cycle progresses (Figure 2C). Therefore, a more complete understanding of the behavioural distribution requires that both the duration and frequency of individual behaviours are examined separately.

**Figure 2.**
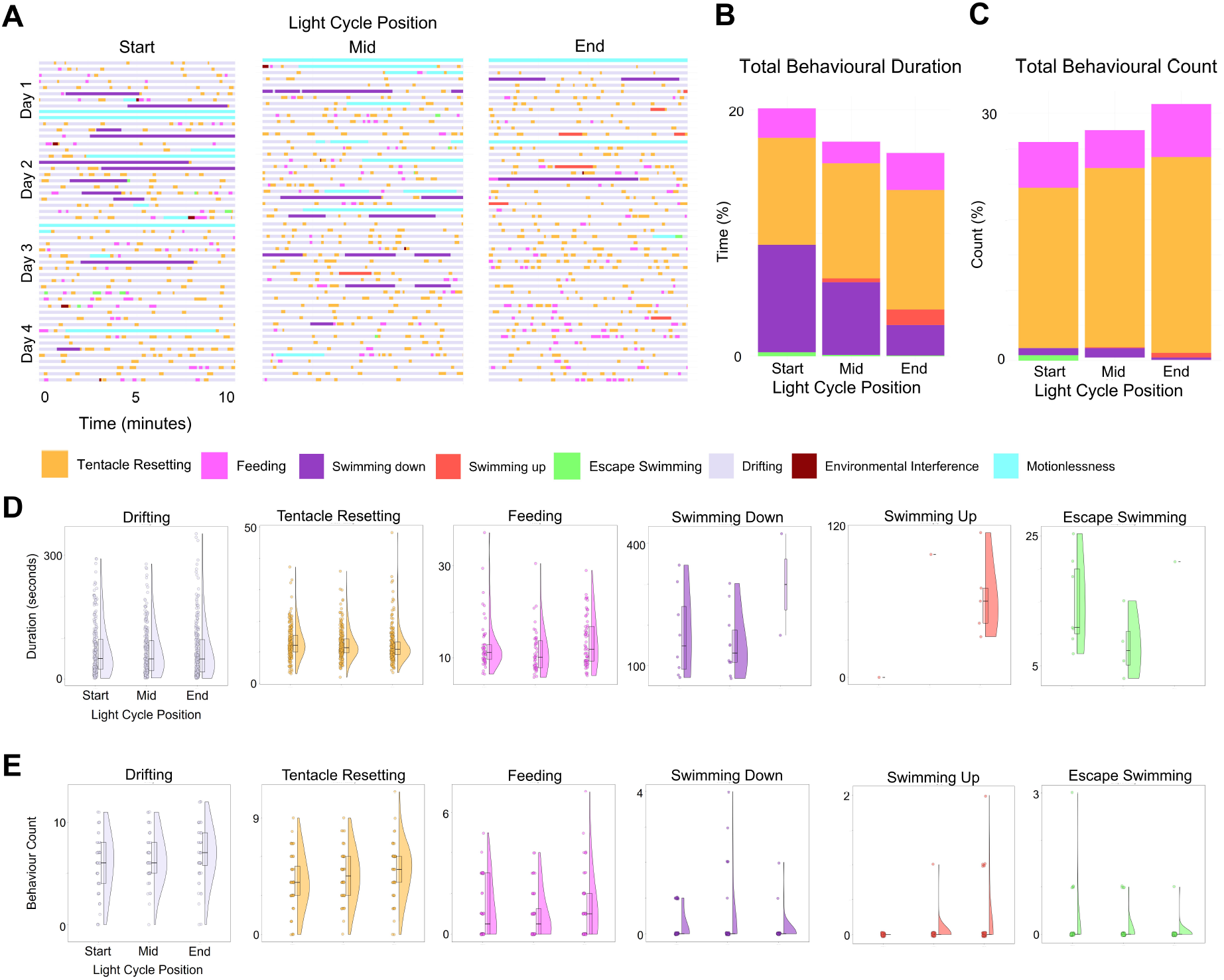
Characterising *Pleurobrachia pileus* Behaviour in Captivity. (A) A timeline of all 10-minute observations at all light cycle positions over four consecutive days. Each behaviour is designated a colour which is used consistently throughout the figures in this manuscript. Active behaviours include tentacle resetting, feeding, swimming up, swimming down and escape swimming. The majority of the time is spent drifting, their default inactive behaviour. Lost data/unclassifiable behaviours due to motionlessness and environmental interference are also noted. (B) The summed duration of each active behaviour at each light cycle position. (C) The summed behaviour counts (number of times a behaviour was initiated) of each of the active behaviours at each light cycle position. (D) The duration of the individual instances of behaviours were extracted at each light cycle position. (E) The number of times a behaviour was initiated for each 10-minute observation was extracted at each light cycle position (behaviour count). In D and E the data is represented as the individual data points, a half violin plot to visualise the distribution and a boxplot which includes the median value as the middle horizontal line and the 25th and 75th quartile range as the lower and upper horizontal lines respectively. The x and y axes labelling conventions used in Tentacle Resetting in D and E are consistent for all plots, however, the y axes limits are different. Note: some behaviours had no instances or few instances in some of the light cycle position groups and therefore do not display a distribution (half violin) or boxplot.

### 3.2 Active Behaviour Duration

We examined whether the duration of a behaviour varied across the light cycle by comparing the durations of all instances of that behaviour across the three time points recorded (Figure 2D). We found no difference between the duration of behaviours at different points in the light cycle for either of the two most common behaviours, namely tentacle resetting (Kruskal-Wallis rank sum test (KW); 2=353.01, p=0.17, df=329, n=671) and feeding behaviour (KW; 2=140.50, p=0.31, df=133, n=168).

Similarly, we found no difference in swimming down across the light cycle (KW; 2=19, p=0.46, df=19, n=20). However, it should be noted that only two instances of this behaviour were recorded at the end of the light cycle. Swimming up was also a relatively rare behaviour with no instances observed at the start of the light cycle position, but no difference in duration was found between the middle and end of the light cycle (KW; 2=5, p=0.42, df=5, n=6). Another rare behaviour was the escape swimming behaviour which had only one instance at the end of the light cycle. Again, no difference was observed between the three time points (KW; 2=11, p=0.44, df=11, n=12). We also tested if drifting duration varied at different times of the day and found no difference (KW; 2 = 680.08, p= 0.39, df=670, n=753).

### 3.3 Active Behaviour Count

While we found no evidence that the duration of instances of active behaviours changed across the light cycle, we also examined whether the frequency of initiation of these behaviours changed over the light cycle. This was assessed by counting the number of times a behaviour was initiated by each animal in a ten-minute recording (Figure 2E). While there was no change in tentacle resetting count across the light cycle (KW; 2=13.73, p=0.19, df=10, n=156) the count of feeding behaviours did show changes, with feeding behaviour initiated more often at the end of the light cycle (KW; 2=15.86, p=0.01, df=6, n=156).

The other three behaviours (swimming up, swimming down, and escape swimming) were, as noted earlier, rarer events. The most common outcome was animals where no incidences of these behaviours were recorded in the 10 minute period of observation. Nonetheless, we noted a difference in the frequency of swimming down behaviours, with higher counts seen at the start of the light cycle (KW; 2=12.72, p=0.01, df=4, n=156), while the opposite was true of swimming up behaviours, which had higher counts at the end of the light cycle (KW; 2=8.06, p=0.02, df=2, n=156), and no instances of swimming up observed at the start of the light cycle. On the other hand, no differences were found between the counts of escape swimming at each light cycle position (KW; 2=3.14, p=0.30, df=2, n=156). When we examined the behaviour count of drifting at different light cycle positions, we also found no difference (KW; 2= 13.83, p=0.31, df=12, n=156).

### 3.4 Examining the Sequential Relationships Between Behaviours

In addition to the frequency and duration of individual behaviours, we also explored how individual behaviours might sequentially influence each other. Specifically, we examined whether individual behaviours were linked into sequential repertoires, which could be identified by repeating sequences of individual behaviours. To do this, we looked at each instance of an active behaviour initiation after a period of drifting (first behaviour), and examined the next behaviour initiated (second behaviour). Two behavioural sequences accounted for the majority of all observations at the start, middle and end of the light cycle: tentacle resetting followed by drifting (Start: 76%, Mid: 78%, End: 78%) and feeding followed by drifting (Start: 21%, Mid: 17%, End: 19%) (Figure 3A). This indicates that active behaviours are not usually chained into sequences, but rather are initiated after a period of drifting, and end with the animal returning to an inactive drifting state before another active behaviour is initiated.

**Figure 3.**
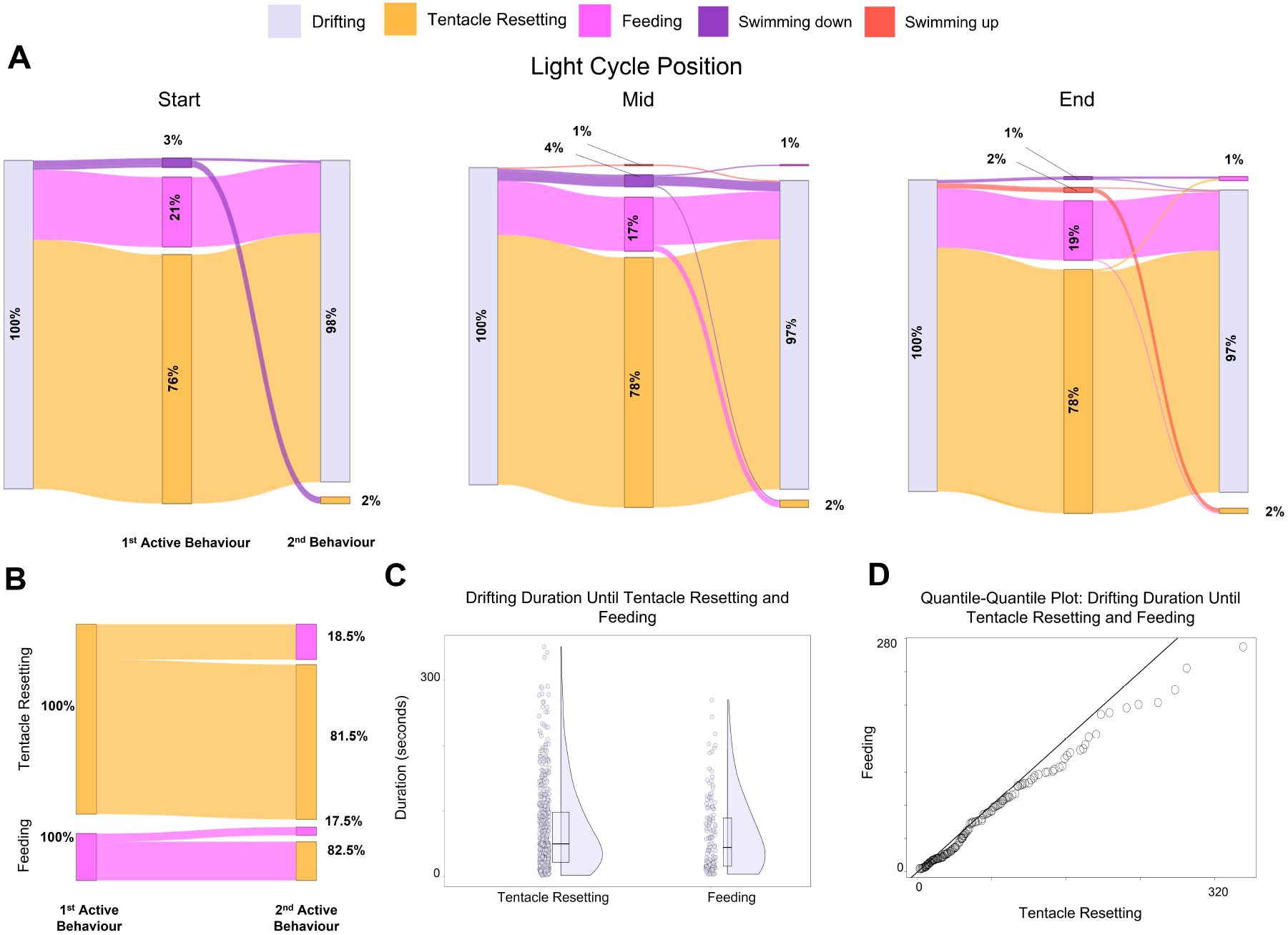
Does a preceding behaviour predict a proceeding behaviour? (A) We extracted all instances of an active behaviour after drifting (1st behaviour) and showed the following behaviour which proceeded it (2nd behaviour). The events are displayed as a percent of all instances in each light cycle position. We next wanted to examine what drives the most common sequence of behaviours and whether one behaviour could predict another. (B) The sequential characteristics of behaviours which begin with a tentacle resetting or feeding behaviour. The events are displayed as a percent of all instances of a sequence beginning with either tentacle resetting or feeding. (C) We pooled all instances of drifting duration until tentacle resetting and feeding across all light cycle positions. This plot includes the individual data points, a half violin plot to visualise the distribution and a boxplot which includes the median value as the middle horizontal line and the 25th and 75th quartile range as the lower and upper horizontal lines respectively. (D) A quantile-quantile plot compares the distributions of drifting interval until tentacle resetting and drifting interval until feeding. No difference between the groups was observed.

While one behaviour did not typically lead directly to the initiation of another behaviour, it remained possible that the initiation of one behaviour could still influence the next active behaviour to be initiated after a period of drifting; specifically, we wanted to determine if the occurrence of either of the two most common active behaviours (feeding or tentacle resetting) predicts the next behaviour initiated after a period of drifting. For example, are animals more like to initiate a tentacle reset after a period of feeding?

Pooled across all observations, resetting behaviours (780 in total) were initiated more commonly than feeding behaviours, (149 in total), giving a percentage ratio for these behaviours of 84%:16%. When feeding was initiated, the next behaviour initiated was also feeding in 17.5% of cases, while it was resetting in 82.5% of cases. When resetting was initiated, the next behaviour initiated was resetting in 81.5% of cases, while it was feeding in 18.5% of cases (Figure 3B). There is no evidence that either behaviour predicts the sequential occurrence of the other (Pearson’s Chi-squared test; 2=1.39, df=2, p=0.50), leaving us to conclude that the behaviours are independent of each other.

### 3.5 What Triggers a Tentacle Resetting Event?

The trigger for the feeding behaviour is well understood: prey captured on a tentacle initiates a cascade of events culminating in prey ingestion. However, the driver of tentacle resetting is less obvious. We decided to examine this question by looking at what leads up to the initiation of tentacle resetting, specifically t he d uration o f drifting until a tentacle resetting event occurred. We can make an assumption that contact with prey, and therefore feeding behaviour initiation, is a random event. Therefore, the interval of inactivity (i.e. drifting duration) before a feeding behaviour should also be random. A behaviour with the same distribution of drifting intervals as feeding is also likely to be random. We then compared the distribution of the drifting duration until tentacle resetting and feeding using a Kolmogorov-Smirnov test. We found no difference between the distribution of drifting durations between feeding and tentacle resetting (Kolmogorov-Smirnov; D=0.11, p=0.12) (Figure 3C-D). Therefore, if the time until feeding is random, it appears that the time until tentacle resetting initiation is also random.

### 3.6 Spacial Characteristics of Feeding and Tentacle Resetting Behaviour Initiation

While analysis comparison of temporal aspects of the initiation of feeding and tentacle resetting behaviours show no apparent differences in their regulation, we must also consider the other external factors. In the aquaculture environment, a key variable may be location within the tank. In the circulating flow of the Kriesel system not all regions of the tank will have the same environmental conditions, particularly in relation to flow. To facilitate this analysis we divided the tank into 5 regions with approximately equals volumes (Figure 4A). The demarcations were decided based on gross fluid dynamics within the tank. Region 1 includes horizontal flow at the top of the tank, region 2 involves downward flow on the right of the tank, region 3 includes horizontal flow at the bottom of the tank, region 4 includes upward flow on the left of the tank and region 5 is in the centre of the tank directly in front of the outflow fi lter. The inflow spray bar was located in region 1 close to the border with region 2. Given the direction of circulation, we assume therefore that region 2 has the most turbulent flow.

**Figure 4.**
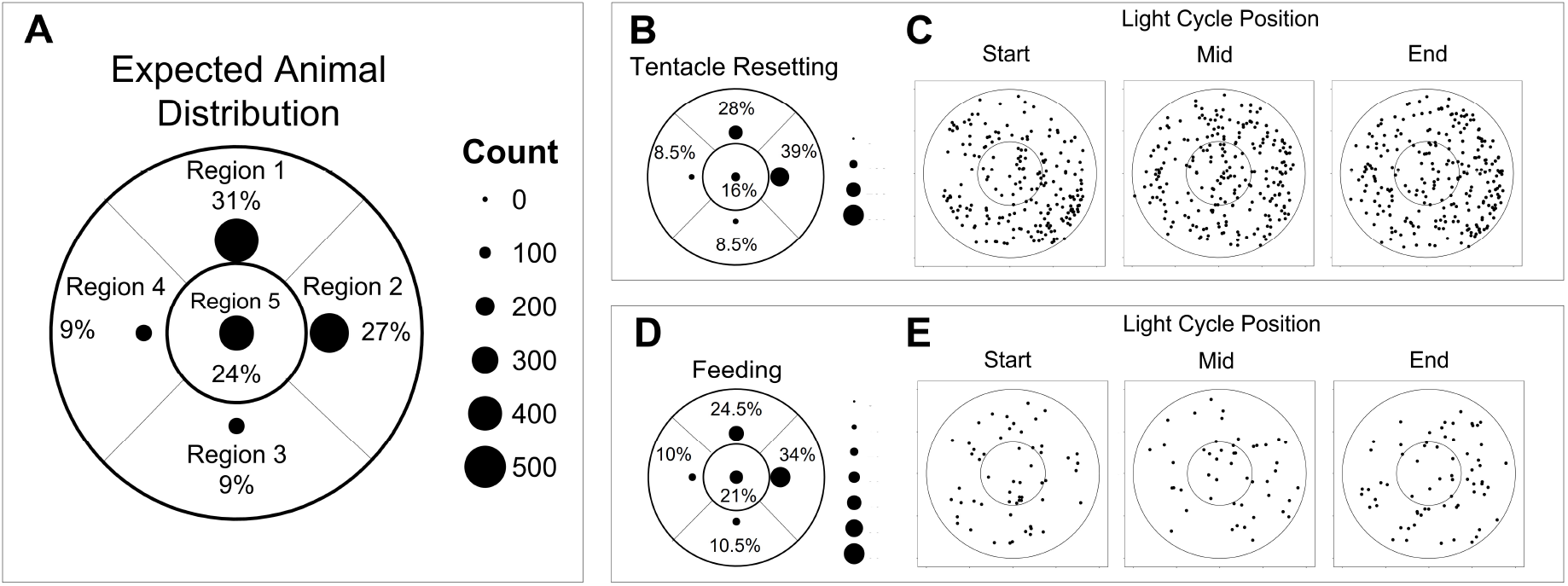
Spatial Characteristics of Behaviours. Does the animal’s location in the tank predict the initiation of a behaviour? We divided the tank into 5 regions, largely based on the assumed fluid dynamics within the tank. We could not assume the animal distribution to be equal across all regions. (A) We calculated the ‘expected’ animal distribution per region by manually counting the location of animals every minute in all videos irrespective of behaviour. (B) The number and proportion of animals which initiated a tentacle resetting behaviour in each region. (C) The location within the kreisel tank upon which tentacle resetting was initiated at each light cycle position. (D) The number and proportion of animals which initiated a feeding behaviour in each region. (E) The location within the kreisel tank upon which feeding was initiated at each light cycle position. A, B D all have the same plotting conventions; the number of instances of an initiation or completion events are displayed as count (point size) and as percent of all instances across all light cycle positions. In all plots the outer circle denotes the outer edge of the tank while the inner circle denotes the edge of the centrally located outflow filter.

We noted the location where each of the feeding and tentacle resetting events were initiated. By comparing the distribution of behavioural initiations across the 5 regions to the distribution of time the animals spent in each region, we could determine if behavioural initiations were more likely to occur in specific r egions. W hile e ach r egion was of the same volume, the time animals spent in each region (the “expected” value for each region) was not equal. The “expected” value was generated by manually counting the distribution of animals every minute in all videos (13 animals at each of the 120 observations) irrespective of behaviour (Figure 4A). We found a significant difference between the distribution of resetting events and overall distribution of the animals (Pearson’s Chi-squared test; 2=38.94, df=4, p<0.0001). It appears that there is a higher probability that animals will initiate tentacle resetting in region 2 (Figure 4B/C). However, we found no difference between the regional distribution of feeding events and the overall distribution of the animals (Pearson’s Chi-squared test; 2=5.52, df=4, p=0.24). It appears the probability of a feeding event occurring in any location in the tank is equal (Figure 4D/E).

## 4 Discussion

We have comprehensively examined the *Pleurobrachia pileus* behavioural repertoire. We analysed the relationships between behaviours, behavioural durations, behavioural frequencies and spatial characteristics of behaviours. This was done to elucidate possible intrinsic and extrinsic mechanisms which drive these actions and also to examine the affect time of day has on them. Of the five active behaviours identified, tentacle resetting and feeding were the most commonly initiated. Swimming down and swimming up were rare but had the longest durations. Behaviour duration does not vary at different times of the day but behaviour count does, for some behaviours. Most of the time was spent in the inactive drifting state. The duration of these periods of inactivity are variable and most often results in transitioning into tentacle resetting or feeding and back to drifting again. The spatial location of an animal in the tank appears to influence the initiation of tentacle resetting.

The most common active behaviour was tentacle resetting. The tentacle resetting behaviour was observed 671 times and accounted for ~9.2% of the total data set. Our data suggests that tentacle resetting is not influenced by a preceding feeding behaviour, it is random in its likelihood of initiation in terms of time but not in terms of its spatial location. This mechanism of preparing the tentacle for prey capture has not been investigated in a detailed manner but has been observed before [Tamm and Moss, 1985]. The aim of this behaviour is presumably to prepare the tentacle for prey capture and performing these motions while relaxing the tentacles and tentilla appear to provide a large surface area. The animals were more likely to initiate a tentacle resetting bout in tank region 2, which due to being downstream of the water inlet is likely to have the most turbulent flow. It seems most likely that this behaviour is not internally generated, but rather is a response to external stimuli, but the nature of the external stimuli are not clear. However, it seems likely that water flow and turbulence, which may impact on the tentacles, is a significant factor in initiating this behaviour.

Feeding and tentacle resetting are closely associated behaviours as they both contribute to successful ingestion of prey. The feeding behaviour in *Pleurobrachia* has been described comprehensively in both free swimming [Tamm and Moss, 1985] and restrained animals [Moss, 1991, Moss and Tamm, 1993]. *Pleurobrachia* feeding behaviour was observed 168 times and accounted for ~2.4% of our total data set. The overall frequency observed in our analysis may be due to many factors; the density of artemia, the kreisel volume and the density and size of *P. pileus* during the time of data collection [Gibbons and Painting, 1992]. It is widely accepted that *Pleurobrachia* is a voracious non-selective predator. Feeding behaviour duration did not vary with the time of day but it did vary in terms of frequency at different time points. This difference may be due to the density of artemia in the tanks. The location of *Pleurobrachia* in the tank does not predict the initiation of a feeding behaviour, which is unsurprising if the prey are distributed in the tanks in a similar manner to the animals. In addition, our results which indicate that a previous tentacle resetting or feeding event did not influence a subsequent feeding event demonstrates the voracious predatory nature of the animals. They do not appear to be using their previous ingestion of prey to prevent a subsequent ingestion. Perhaps at higher food availability their satiety would impact the initiation of a feeding behaviour. However, we have witnessed animals purging gut material and this may be the only mechanism they have to control their food intake. While tentacle resetting and feeding were the dominant active behaviours, we also observed other distinct behaviours that, while rarer, may also play important roles in ctenophore biology, although the actual function of the behaviours is less clear. Swimming up and down are two behaviours which fall into this category. Animals which demonstrated swimming up and swimming down behaviours had to be undertaking this act for more than 28 seconds to be constituted as such because feeding and tentacle resetting were also associated with preceding directional swimming. These behaviours are the longest duration active behaviours, but they are not very common. Six instances of swimming up and twenty instances of swimming down were noted from the entire data set. In all cases of swimming up, it exclusively occurred on the right side of the tank as the animals swam against the water flow. In contrast, swimming down exclusively occurred on the left side of the tank where they again were swimming against the water current. Swimming up equates to ~0.5% of the total data set while swimming down equates to ~6%. However, we must consider the impact the tank walls will have on these behaviours. Perhaps the durations and counts would be higher in the wild with less obstacles disturbing the response. Encountering the tank walls during this behaviour may cause shorter durations. The aquaculture system we describe here may not be ideal for studying these specific behaviours. We observed that the duration of the swimming up and down events did not vary with the time of day but the behaviour counts did. While it is tempting to link this observation to possible diurnal migration, we must be cautious in drawing conclusions from relatively rare behaviours.

The behaviours which we referred to as escape swimming we believed encompassed high frequency active swimming behaviours that did not fall into the realm of the other active behaviours and that were more ‘erratic’ in nature. They are relatively rare (12 instances and ~0.2% of total data set). There was no difference in duration or count of this behaviour at any of our sampled time points. However, we did observe that 83.5% of initiation events occurred in region 5. Region 5 encompasses the filter outflow; perhaps body wall or tentacle interactions with a filter pore may have driven this behaviour. Our classification of an ‘escape’ behaviour appears to closely resemble behaviours in the literature. Global ciliary excitation is described as a fast forward swimming response [Tamm, 1982]. Global ciliary excitation can also occur in reverse [Tamm and Tamm, 1981, Mackie et al., 1992, Tamm and Terasaki, 1994]. Our results indicated that there isn’t a substantial difference in behavioural characteristics at different times of the day. Our analyses may have missed specific features of a circadian rhythm, or the aquarium environment may have altered their natural circadian rhythms, so we cannot rule out the existence of a circadian rhythm in these animals. Circadian rhythms are influenced by light, but direct physiological evidence of light-sensing cells in *Pleurobrachia* does not exist. Chun [1880], Horridge [1964] and Hernandez-Nicaise [1984] described structures resembling photoreceptors in the aboral organ. In addition, a more recent investigation in *Mnemiopsis leidyi* found opsin expression co-localised with these structures which supports their role as photoreceptors [Schnitzler et al., 2012]. Ctenophores have been shown to respond to light behaviourally; spawning [Freeman and Reynolds, 1973, Pianka, 1974, Pang and Martindale, 2008, Patry L. et al., 2019] and reversing geotaxis [Tamm, 1982].

This list of behaviours in our analysis is not exhaustive, due to the nature of our data collection and analyses there were certain behaviours we could not identify and may have been included under the umbrella of others. For ex ample, mechanical stimulation to any location on *Pleurobrachia’s* body results in temporary interruption of beating of all the comb plates, this has been referred to as sudden stoppage/global ciliary inhibition/primary inhibition [Moss, 2004]. Another response not noted in our data set is secondary inhibition, stronger stimulation (as compared to primary inhibition) leads to muscular contraction of the comb rows inward presumably to protect the important locomotory structures [Tamm and Tamm, 1989]. Subtler actions such as body wall muscular contractions, mouth movements, defaecation [Presnell et al., 2016] and gamete release [Pianka, 1974, Patry L. et al., 2019] could also not be captured in this analysis.

Our results also show that collecting a sufficient number of animals is feasible and reliable at Howth Harbour, but their density is seasonal. Changes in overall density of *Pleurobrachia* at different times of the year and between years has been observed by many researchers [Hirota, 1974, Yip, 1984, Frank, 1986, Buecher and Gasser, 1998]. The seasonal peaks of *P. pileus* in different locations was summarised by Wang et al. [1995] with the highest densities for most reported in May and June. This seasonality makes a long-term aquaculture system necessary for reliable, year-round experimentation. Our aim was not to develop an optimal aquaculture system, but to develop one that was simple, robust and sufficient for our purposes, specifically providing long term reliable access to animals without necessitating on demand collection at sea, and allowing observation of animals for a period of several weeks at least. The survivability data is challenging to interpret as the life span of *P. pileus* in the wild is unknown. Factors which may have impacted the longterm survival and behaviour of these animals includes animal density, food source, temperature, lighting, light-dark cycle and water quality. Potential improvements include rigorous monitoring of environmental variables and assessing the impact of animal density and food sources on survival and behaviour. Future studies could take advantage of a deep neural network approach to tracking animal trajectory [Mathiset al., 2018] and the development of a machine learning algorithm could also enable an unbiased method of categorising behaviours [Han et al., 2018]. These tools would enable longterm and high-throughput behavioural analysis. A successful ctenophore culture protocol has recently been established at the Monterey Bay Aquarium [Patry L. et al., 2019] which could easily be combined with the system we described here. However, many benefits exist to studying wild caught animals it indicates that their nervous system was working optimally to enable them to develop, avoid predators and acquire food.

## Conclusion

We have shown that *P. pileus* can be collected reliably and in high densities at specific times of the year in the Irish Sea and that we can keep them alive in a specialised aquarium system for months at a time. We hope the description of our ctenophore husbandry system will encourage others to publish detailed accounts of their efforts in this regard.

The optimisation and standardisation of these systems are imperative to the establishment and longevity of a successful ctenophore research community. We also characterised the full behavioural repertoire of *P. pileus* in our aquarium system. This analysis may be of interest to neuroscientists who are interested in answering questions about the foundational principles of nervous systems in a tractable model. It is unclear if our behavioural profile is comparable to their behaviour in the wild and an important next step will be to acquire a similar dataset from animals in their natural environment.

## Acknowledgements

The authors wish to acknowledge Dominic Courtney for his invaluable assistance in the collection of ctenophores. The authors would also like to thank the staff at Seahorse Aquariums for their expert advice in setting up the aquaculture system.

## Statement of Ethics

As they are invertebrate species, use of ctenophores in research is exempt from full ethical review under local policies and legislation. Formal exemption from review was granted by UCD AREC (AREC-E-17-22-Pickering).

## Author Contributions

AC collected the ctenophores, designed and built the husbandry system, carried out and analysed the behavioural and survival experiments and wrote the manuscript. GM assisted in the collection and maintenance of ctenophores and designed the automated brine shrimp feeding system. MP conceived the project, supervised all aspects of the work and edited the manuscript.

