## Supplementary figures and images for "Characterising the Behaviour of the Ctenophore *Pleurobrachia pileus* in a Laboratory Aquaculture System"

### Supplementary Table 1

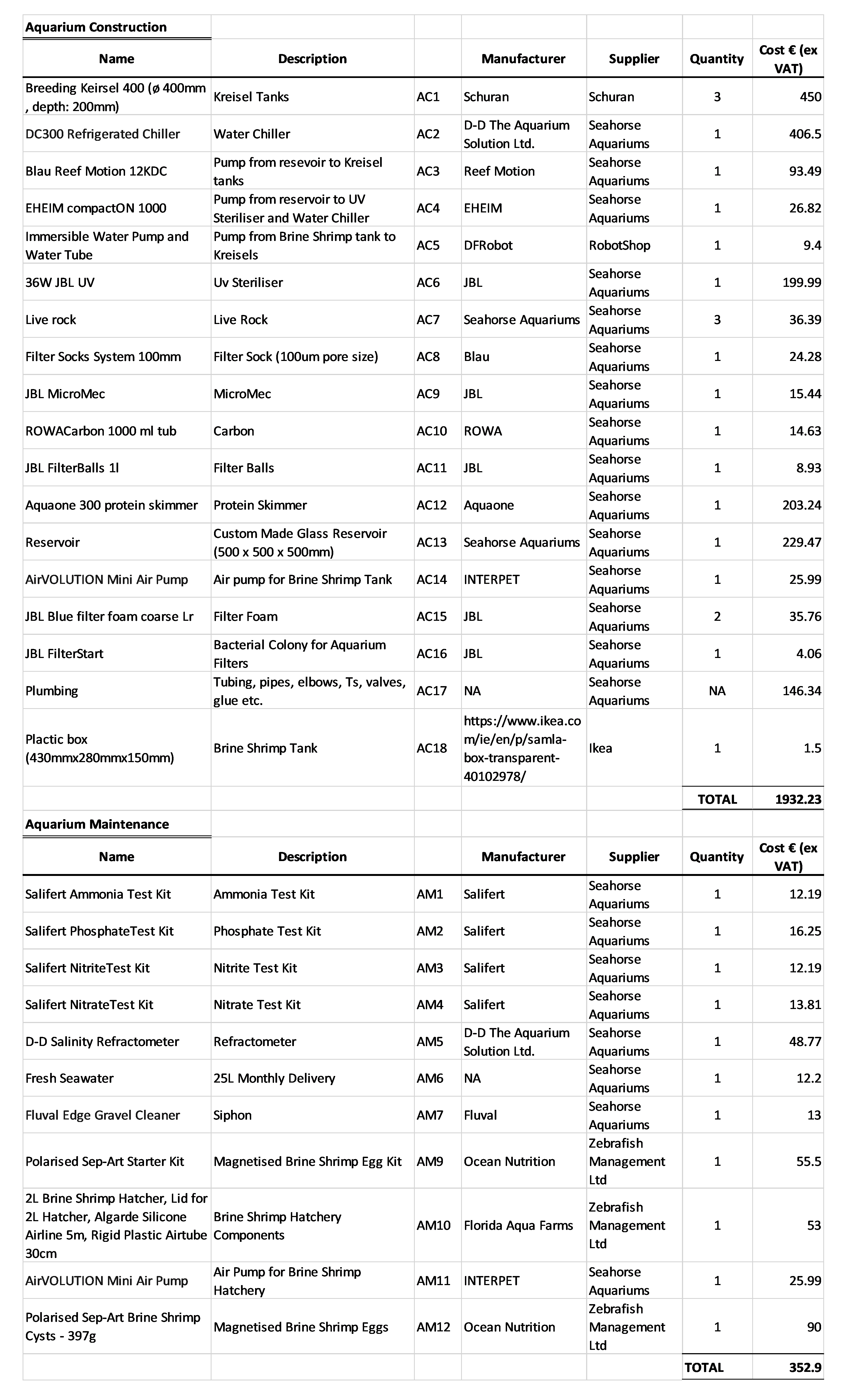

### Supplementary Table 2

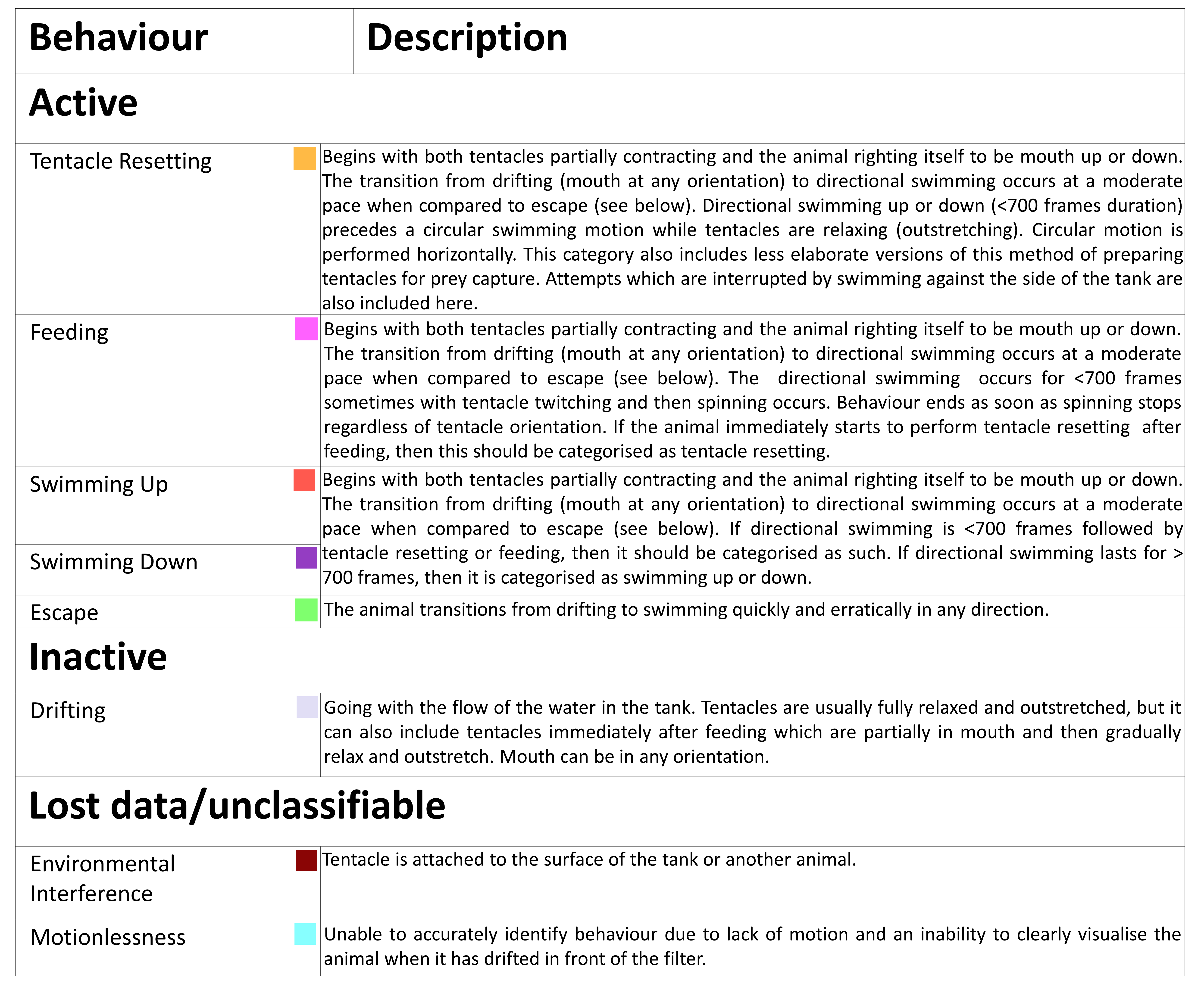
